# Training-induced prefrontal neuronal changes transfer between tasks

**DOI:** 10.1101/2020.10.22.351197

**Authors:** Hua Tang, Mitchell R. Riley, Balbir Singh, Xue-Lian Qi, David T. Blake, Christos Constantinidis

**Affiliations:** Department of Neurobiology & Anatomy, Wake Forest School of Medicine, Winston-Salem, NC 27157, USA; Center for Neuropsychiatric Diseases, Institute of Life Science, Nanchang University, Nanchang, Jiangxi 330031, China; National Institutes of Mental Health, NIH, Bethesda, MD 20892, USA; Department of Psychology, Vanderbilt University, Nashville, TN 37240, USA; Department of Neuroscience and Regenerative Medicine, Medical College of Georgia, Augusta University, Augusta, GA 30912, USA

## Abstract

Training to improve working memory is associated with changes in prefrontal activation and confers lasting benefits, some of which generalize to untrained tasks, though the issue remains contentious and the neural substrate underlying such transfer are unknown. To assess how neural activity changes induced by training transfer across tasks, we recorded single units, multi-unit activity (MUA) and local field potentials (LFP) with chronic electrode arrays implanted in the prefrontal cortex of two monkeys, as they were trained to perform cognitive tasks. Mastering different tasks was associated with distinct changes in neural activity, which included redistribution of power across frequency bands in the LFP, recruitment of larger numbers of MUA sites, and increase or decrease of mean neural activity across single units. In every training phase, changes induced by the actively learned task transferred to an untrained control task, which remained the same across the training period. The results explicate the neural basis through which training can transfer across cognitive tasks.

## Main

Working memory, the ability to retain and manipulate information over a period of seconds, represents a core component of higher cognitive functions, including control of attention, non-verbal reasoning, and academic performance ^1–3^. Working memory ability has been traditionally thought of as an immutable aptitude, but it is now understood that it can be improved by training in working memory tasks ^4–6^. The extent over which performance improvement after working memory training generalizes, or transfers, to tasks that were not part of the training has been a matter of debate; some studies have been successful in inducing transfer from one task to another whereas others have not ^4–11^. Less contested is the idea that working memory training is beneficial for patients with clinical conditions, including attention deficit hyperactivity disorder (ADHD), traumatic brain injury, and schizophrenia ^4, 12, 13^.

The neural basis of transfer has been poorly understood. Human fMRI studies have produced conflicting results about the effects of cognitive training, suggesting overall increases ^13–18^, or decreases in activity ^19–22^, or more subtle differences such as changes in network modularity ^23, 24^. Increases are interpreted as reflecting a higher level of activation or recruitment of a larger cortical area, decreases as suggestive of improvements in efficiency ^25, 26^. What these correspond to at the level of neural spiking activity and how lasting changes can transfer between tasks has remained hitherto unexplained. We were thus motivated to address the neural effects of working memory training that could transfer between tasks with neurophysiological recordings in monkeys. Persistent discharges that continue to represent stimulus properties are thought to underlie working memory, though this is a topic of recent debate, as well ^27,28^. We tracked neuronal activity with a chronically implanted electrode array throughout several months of training, and were thus able to address changes in neuronal activity with training and possible transfer between tasks.

## Results

### Monkeys acquire different elements of cognitive tasks with training

Two male Rhesus monkeys (*Macaca mulatta*) were initially acclimated with the laboratory and trained to maintain fixation and not respond to stimuli presented on a computer screen. The monkeys were then trained to perform a spatial working memory task, requiring them to maintain fixation, observe two stimuli appearing in sequence separated by delay periods, and to indicate if the two stimuli appeared at the same location or not by selecting one of two choice targets, defined by their shape (“H” or “Diamond” in Fig. 1a-e). The training required to acquire and master this task consisted of four phases. First, the monkey was presented with two stimuli in rapid succession and had to indicate if they appeared at the same or different locations by selecting one of two choice targets signifying match or nonmatch (Fig. 1b). During this phase, daily sessions involved presentation of the cue at the right of the fixation point followed by a sample stimulus appearing at either a matching location (right) or a nonmatching location (left), on different days. At this stage, the monkey could simply sample the match or the nonmatch choice targets, determine which one was rewarded during the block, and repeatedly select it in following trials. In the second phase, the monkey was presented with alternating blocks of match and nonmatch trials, of decreasing block length, until they were randomly interleaved, requiring the monkey to associate the match and nonmatch conditions with the corresponding choice target (Fig. 1c). In the third phase, the monkey had to generalize the task to new stimulus locations, appearing at a 3 × 3 grid (Fig. 1d). Finally, an increasing delay period was imposed, placing more demand on working memory (Fig. 1e).

**Figure 1.**
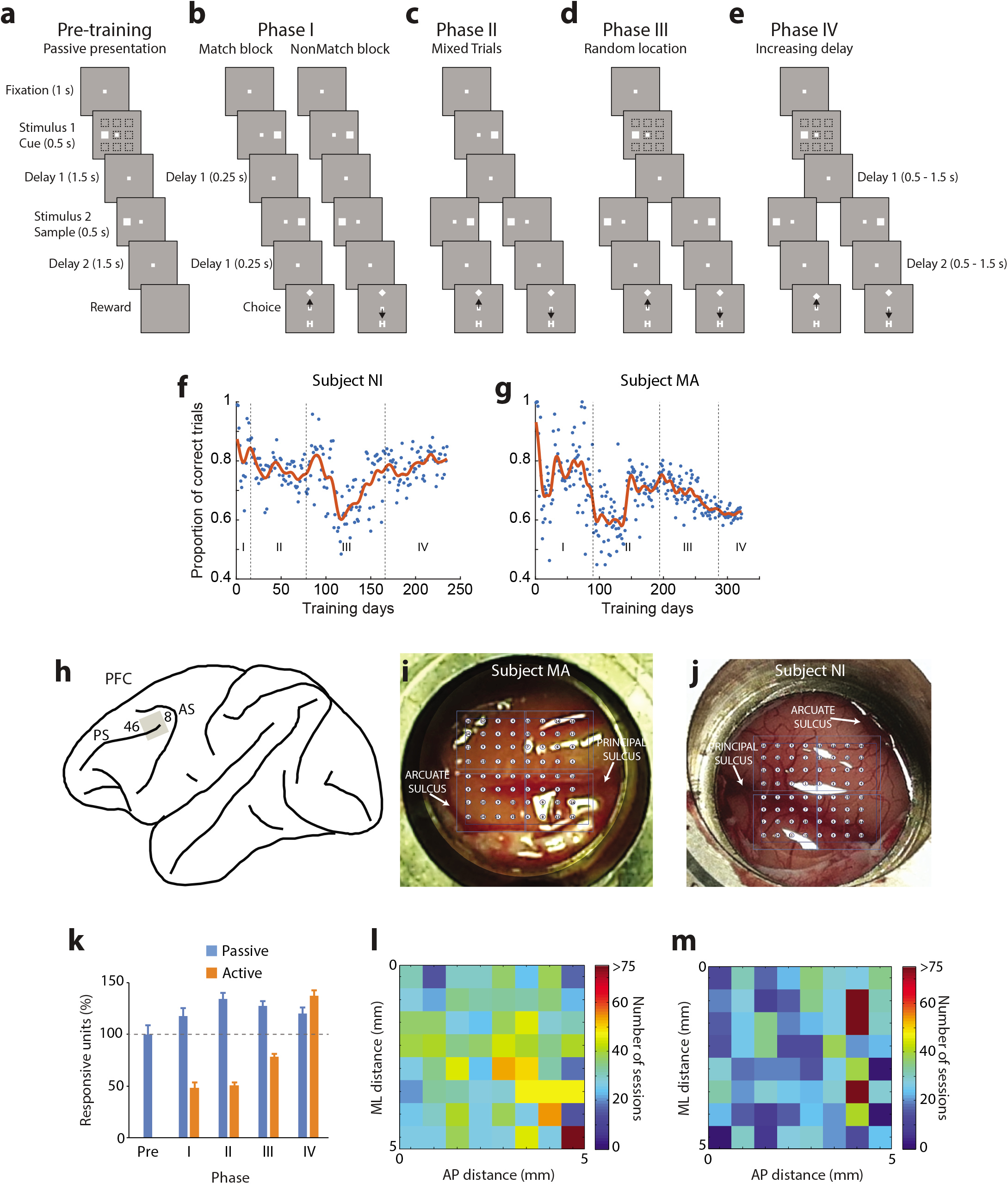
Behavioral Training and Chronic Array. **(a-e)** Successive frames illustrate the sequence of events in the tasks used in progressing training phases. **(a)** During the pre-training phase, the monkey had to only fixate while the stimuli were displayed at any of nine locations on the screen. **(b)** In Phase I, a stimulus was always presented to the right, followed by a match stimulus in a block of trials and by a nonmatch stimulus in another block of trials. At the end of the trial, two choice targets appeared, and the monkey had to choose the “Diamond” target in match blocks and the “H” target in nonmatch blocks to get a reward. **(c)** In Phase II, match and nonmatch trials were mixed in a block. **(d)** In Phase III, the stimulus location of the first stimulus could vary. **(e)** In Phase IV, the duration of the delay period increased. The passive stimulus set continued to be presented at the beginning of each session throughout training. **(f-g)** Performance of two monkeys at each daily session. **(h)** Schematic diagram of the monkey brain with the approximate location of the recording grid (gray square) indicated relative to prefrontal landmarks: areas 46 and 8, Principal Sulcus (PS) and Arcuate Sulcus (AS). **(i)** Position of the electrode array in the right prefrontal cortex of monkey MA is indicated relative to the PS and AS. **(j)** Position of the electrode array in the left hemisphere of monkey NI. **(k)** Relative numbers of responsive units in each training phase for passive and active tasks. The number of units is shown as a proportion relative to the average unit number of the passive task in the pre-training phase. Data from two subjects, for MA, n = 1341 in the passive task, n = 1150 in the active task; for NI, n = 816 in the passive task, n = 1230 in the active task.

Importantly, visual stimuli were also presented to the monkeys passively, in a control fixation task every day (Fig. 1a). The sequence of events in the passive trials mirrored the final phase of the active task; two stimuli were presented at random locations with the second stimulus appearing either at a matching or a nonmatching location, separated by 1.5 s delay periods. The critical difference was that no choice targets were presented in the passive task, and the monkey was rewarded at the end of the second delay period for maintaining fixation and omitting responses to stimuli. The monkeys performed this passive fixation task before recordings began, and they continued to perform it in exactly the same fashion at the beginning of each daily session before active task training began. Training proceeded in an adaptive manner, so that the task became progressively harder as the monkeys mastered each element of the task so that overall performance remained approximately constant through the duration of the training (Fig. 1f-g).

### Training increases neuronal activation and decreases beta power

After initial acclimation with the laboratory, and before Phase I training began, the animals were implanted with a chronic array of electrodes in their lateral prefrontal cortex (Fig. 1h). The implant comprised an 8 × 8 grid of electrodes, with adjacent electrodes spaced 0.75 mm apart from each other, thus covering an area of 5.25 mm × 5.25 mm. The electrode array was implanted in the dorsolateral prefrontal cortex (dlPFC), with electrode tracks descending in both banks of the principal sulcus (Fig. 1i-j). Local Field Potentials (LFP) and Multi-unit activity (MUA) was recorded from all electrodes, which remained fixed after training began. To sample spiking activity in an unbiased fashion, we set the exact same MUA threshold criterion for all electrodes and sessions, to 3.5 × root mean square (RMS) of the noise level. We were thus able to quantify systematic changes in neural activity as training took place. We identified MUA units with responsiveness to stimuli as those exhibiting a significant elevation of firing rate during either the first stimulus presentation or the delay period following it (see Methods). A total of 4537 responsive MUA units were identified in this fashion across all phases of training and across all electrodes, with a sustained yield of responsive units through the last training phase (Fig. 1k-m). Single neuron recordings were also obtained, after spike sorting of the MUA records. We identified a total of 1207 single units responsive to the active task and 1065 responsive to the passive task, based on the same criteria.

We first examined LFP power spectra, averaged across all electrodes and available sessions, which provided an overview of neuronal changes across training phases. Theoretical and experimental studies suggest that improved working memory maintenance is associated with decreased power in the beta-frequency band and increased power in the gamma band ^29–32^. We therefore wished to test the hypothesis that training would produce overall decreases in beta power and increases in gamma. Certain features were present across all phases, such as a broadband power elevation during the appearance of the fixation point (time point “-1” s) in Fig. 2a), a narrower power increase during the cue appearance (time point “0” in Fig. 2a), and a power increase at the time of the saccade and subsequent reward (after the last vertical line in each plot of Fig. 2a). In partial agreement with our hypothesis, training induced systematic changes in power, the most salient of which was a progressive decrease in power in the beta/low-gamma frequency zone of 20-45 Hz (hereafter referred to as beta, for simplicity) during the cue presentation period in successive active training phases (Fig. 2c). Averaging beta power over the entire cue period revealed a highly significant difference between phases (1-way ANOVA comparing beta power in daily sessions grouped in four training phases, F_3,889_ = 113.8, p = 2.27 × 10^-62^). A concomitant increase in alpha-frequency power (8-14 Hz) was also observed (F_3,889_ = 94.0, p = 7.47 × 10^-53^). High gamma (46-70 Hz) power was less diagnostic of the training progression but generally moved in the opposite direction than our initial hypothesis. Importantly, those global changes in beta and alpha power were also present in the passive-fixation task (Fig. 2d), which the monkeys continued to be exposed daily, at the beginning of each session before training in the active task began. Although the passive task stimuli never changed, we observed a significant decrease in beta power across successive stages, considering the pre-training phase as well (1-way ANOVA, F_4,446_ = 33.8, p = 1.28 × 10^-24^), and a relative increase in alpha power (F_4,446_ = 18.0, p = 1.02 × 10^-13^). The decrease of beta power/increase of alpha power across training phases that transferred into the passive task was observed in both monkeys (Fig. S1). The effects were essentially identical when we performed LFP analysis only in electrodes from which single neurons were recorded, to ensure that changes detected were not the result of some electrodes becoming inactive (Fig. S2).

**Figure 2.**
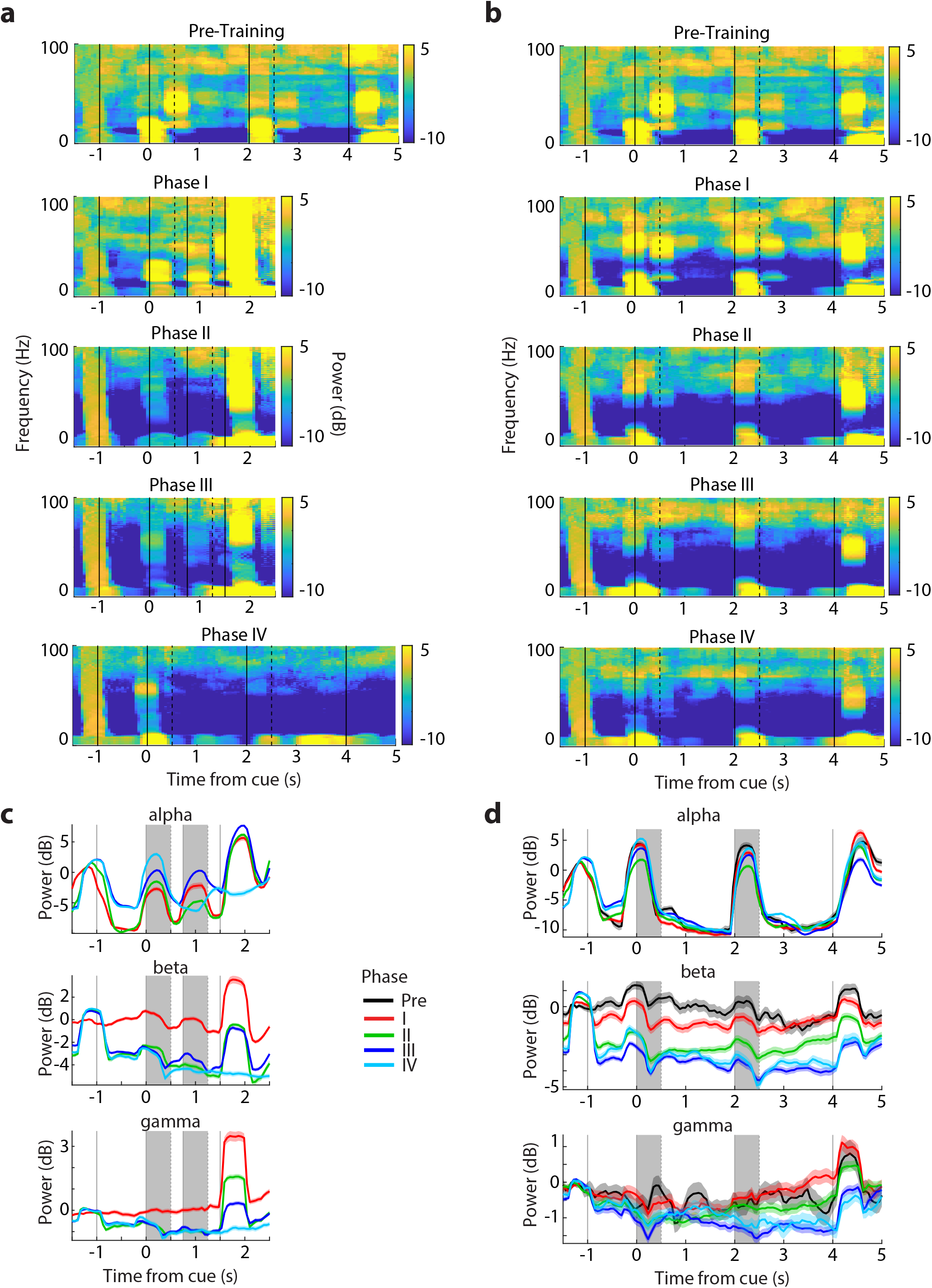
LFP analysis. **(a)** LFP power spectrum as a function of time for the training phases in the active task. The pre-training phase is provided for comparison. Power (μV^2^/Hz) is expressed as a ratio over average power across the spectrum, in logarithmic units. **(b)** LFP power spectrum as a function of time for the passive task, as training progressed in the active task. The pre-training phase is also provided for comparison and is the same as in panel a. **(c)** Time course of power at discrete frequency bands and different training phases of the active task: alpha (8-14 Hz), beta (20-45 Hz), gamma (46-70 Hz). **(d)** Time course of power in the same frequency bands for the passive task, as training progressed in the passive task. Shaded areas represent the stimulus presentation periods.

We next addressed the effects of training on neural activity. Based on experimental and theoretical grounds ^33^ we hypothesized that a great proportion of neurons would be activated, and at a higher firing rate. Indeed, training in the active task resulted in a greater population of prefrontal MUAs becoming responsive to the stimuli (Fig. 1k, orange bars), and in a higher mean firing rate generated by single neurons (Fig. 3a-c). Comparison of mean firing rates for the best location of each single neuron in each training phase, after subtracting baseline activity revealed a highly significant difference between stages (1-way ANOVA test, F_4,807_ = 83.15, p = 4.0 × 10^-59^ for the cue period, F_4,807_ = 20.08, p = 8.93 × 10^-16^ for the delay period). These changes in firing rate were also evident in the context of the passive fixation task (Fig. 3d-f), though changes were not always monotonic or as consistent. Firing rates for the best location after subtracting the baseline was significantly different between phases (1-way ANOVA test, F_4,285_ = 4.1, p = 0.003 for the cue period, F_4, 285_ = 5.41, p = 0.0003 for the delay period). Even though stimuli were presented exactly in the same fashion every day, prefrontal single neurons generated higher levels of activity after they had been trained to perform a task.

**Figure 3.**
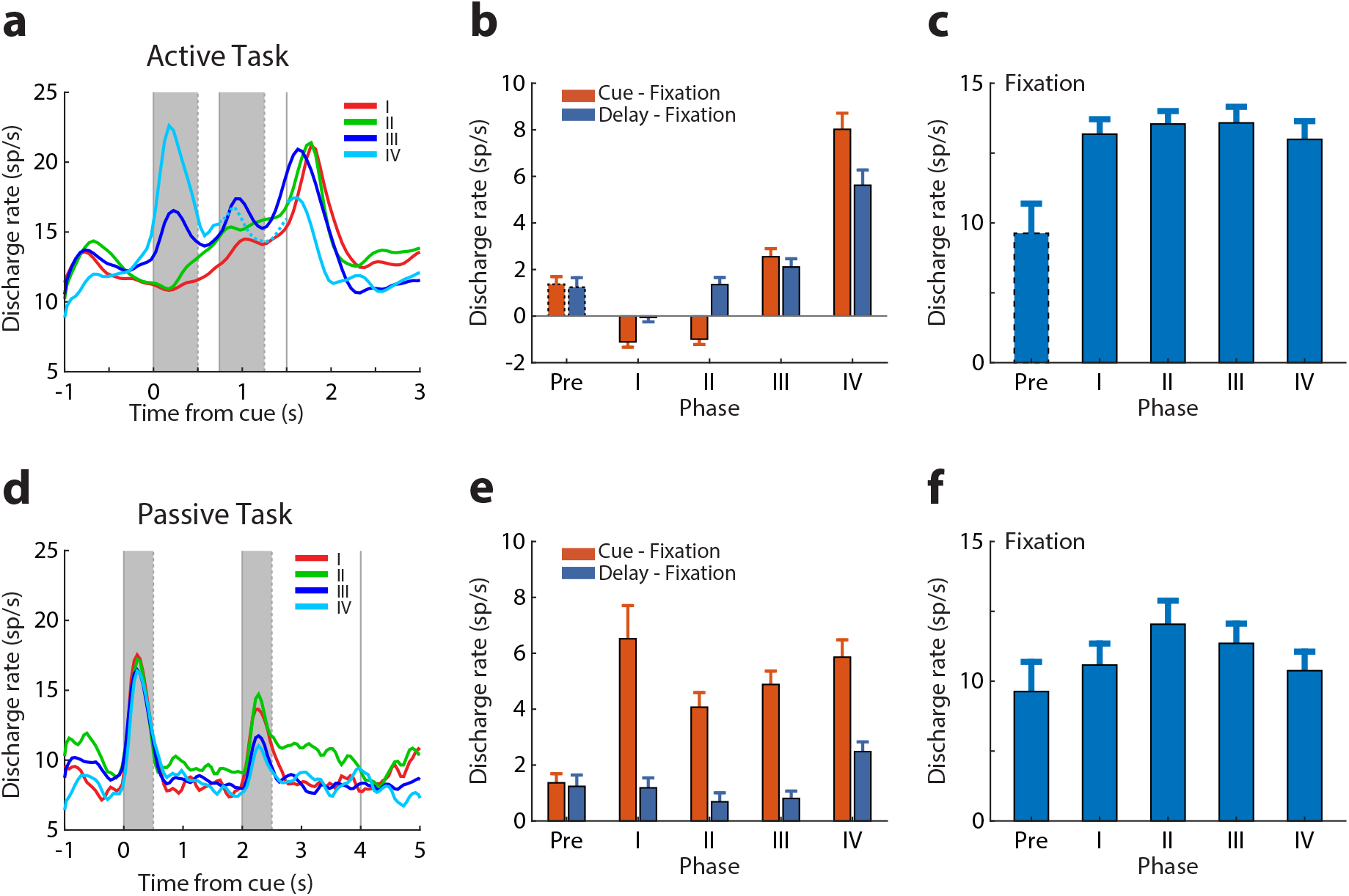
Mean firing rate of single neurons at different training phases. **(a)** Population peri-stimulus time histogram (PSTH) of responsive neurons in the active task (n = 1207). Best stimulus location for each responsive neuron is used, aligned to the cue presentation. Shaded areas represent the stimulus presentation periods. The delay period was variable in phase IV; only the first 250 ms are indicated (activity followed the second stimulus is plotted in dotted line); the rest of the plot is aligned to the response onset. **(b)** Neuronal activity averaged over the cue and delay periods after subtracting the baseline is plotted for each of the training phases. **(c)** Baseline fixation for the active task. **(d)** Population PSTH of all responsive units in the passive task (n = 1065). **(e-f)** Data plotted as in panels b and c, for the passive task.

The cumulative effect of a greater population of units being recruited and firing at a higher rate during the passive task as training in the active task progressed could be appreciated when we tracked MUA activity from the same channel over repeated days. Absolute activity in the example channels illustrated in Fig. S3 peaked at stage III (when the monkey mastered the full task, in active training sessions practiced later in the day). This increase in firing rate was evident already from the baseline fixation interval, though peak cue and delay period activities also changed during the course of training. It was also important to realize that the firing rate changed continually even within each phase, as the monkey figured out new elements of the task and improved in performance. This can be appreciated when we plotted the MUA firing rate on a day-to-day basis, as training progressed (Fig. 4a-c). This illustration also made evident that a more granular analysis was necessary to understand the nature of neuronal activity changes during training and how these transferred between tasks.

**Figure 4.**
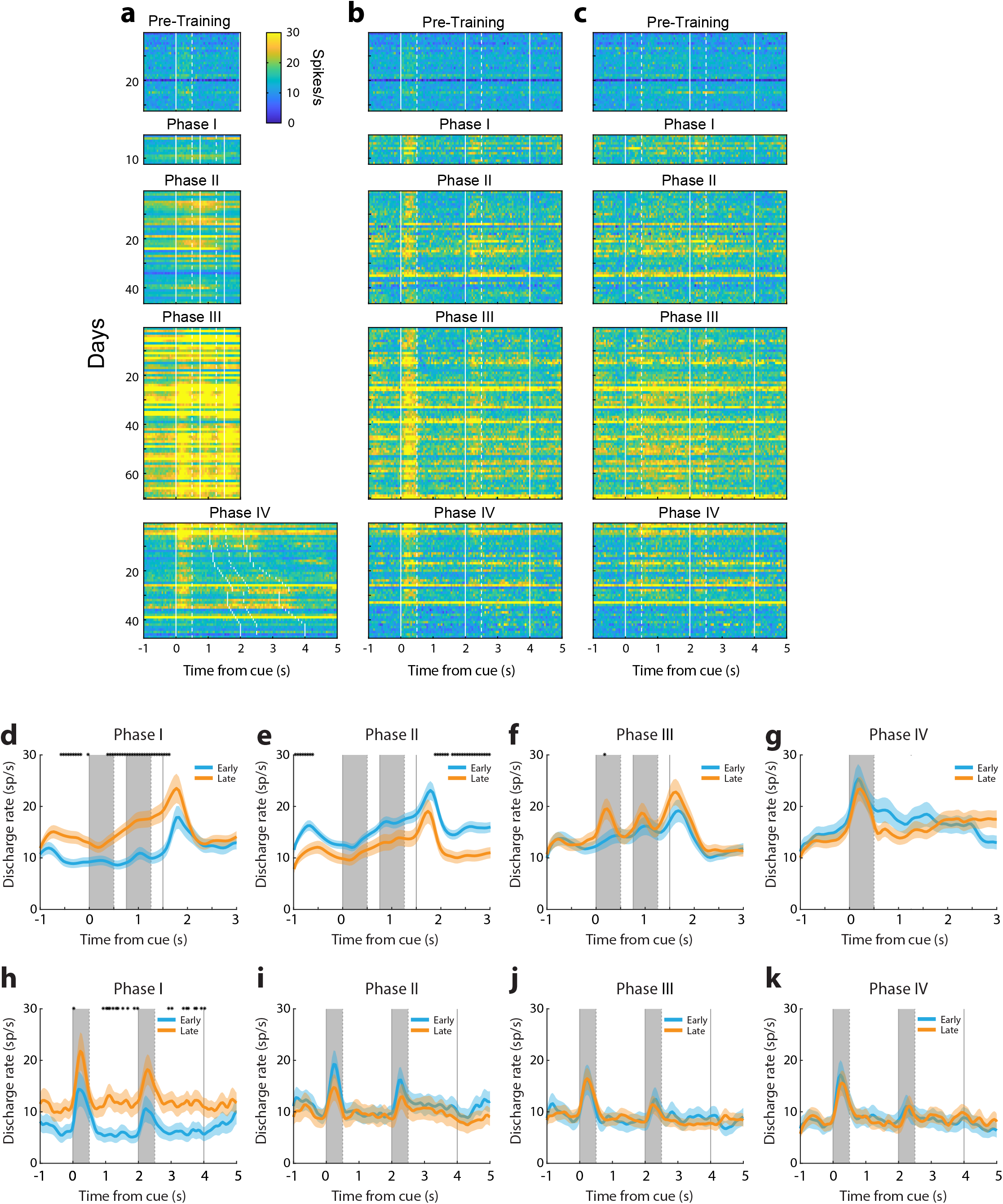
Daily responses in the active and passive tasks as the active training progressed. **(a-c)** Activity of MUA units responsive to the active task (a) and the passive task (b-c). Color plot represents the mean firing rate of all responsive MUA units available on that day. Only days with responsive MUA units in both the active and passive tasks were identified are shown. Data are plotted for the best cue location (a-b) and the best delay period (c) activity of the MUA units under study. **(d-g)** Population PSTH of responsive neurons in the active task (n = 1207). **(h-k)** Population PSTH of responsive neurons in the passive task (n = 1065).

### Neural effects of acquisition of different task element transfer between tasks

Training in Phase I required the monkeys for the first time to observe the choice targets and select one as a saccade target (Fig. 1b), creating associations between sensory stimuli and reward or its omission. We point out that in the pre-training phase, if a monkey responded to any stimulus, the reward was omitted. Trials with cue and match presentations alternated with cue and nonmatch presentations, in different sessions. The subject could perform the task by simply ignoring the two first stimulus presentations, waiting until the choice targets appeared, and testing which one of the two was rewarded, then returning to the rewarded target in all subsequent trials of the session. We hypothesized that the significance of these task events would be reflected in neural variables. Indeed, a broadband power peak in the LFP (Fig. 2a) and a peak in firing rate (Fig. 3a) was evident at the time the choice targets appeared. Little phasic response was evident during the presentation of the cue and match/nonmatch.

However, activity ramped during the time course of the trial, peaking before the appearance of the choice targets (Fig. 3a). We also postulated that the active engagement in the task would result in heightened activation of the prefrontal cortex in the baseline period and during the presentation of the visual stimuli, now in the context of a task. This expectation was also confirmed. During the course of learning the association between choice targets and reward, long-lasting changes in the prefrontal network were observed, which also transferred during the passive task: firing rate during the execution of the active task increased during the course of training (Fig. 4d, twotailed t-test, t (134) = 3.07, p = 0.003). The same rate change was also observed in the passive task (Fig. 4h, two-tailed t-test, t (39) = 2.22, p = 0.032).

In Phase II, presentations of both match and nonmatch trials occurred during the same session. At the initial training sessions, match trials were presented until the subject completed 50 correct responses, and these were followed by nonmatch trials. In this stage, too, the subject could perform the task by ignoring the cue and match/nonmatch stimulus and relying on reversal learning of the rewarded choice target. However, as the blocks of match and nonmatch trials became shorter, and eventually fully randomized, the subject could only perform the task by becoming aware that the “Diamond” choice target was associated with the match stimulus and the “H” choice target with the nonmatch (Fig. 1c). This training introduced a new type of association between reward and a cognitive abstraction, the concept of “match” and “nonmatch”. We note that throughout stage II, the monkeys could still perform the task by essentially ignoring the cue (first stimulus), since it always appeared at the same location. We hypothesized that the significance of these task events would also be reflected in neural variables. Indeed, little response continued to be present during the cue period (Fig. 3a-b), but the firing rate further accelerated during the second stimulus presentation, prior to the saccade (Fig. 3a). Based on experimental results in the sensory cortex, we hypothesized that this training would induce a transient, non-selective increase in responsiveness ^34^, which we expected would be reflected in baseline and stimulus-driven firing rate. We indeed observed changes in the prefrontal network, which also transferred during the passive task: firing rate during the execution of the active task initially increased relative to Phase I, but then decreased again during the course of training (Fig. 4e, two-tailed t-test, t (298) = 2.12, p = 0.035). A parallel pattern of rate changes was not observed in the passive task (Fig. 4i, two-tailed t-test, t (65) = 0.49, p = 0.624). These effects were evident in the day-to-day changes (Fig. 4a-c). We note that different neurons were responsive in the active and passive task; these changes reflected overall changes in responsiveness across the prefrontal network, rather than sampling of neurons with lower or higher activity at different recording dates.

Training in Phase III required the subjects to generalize across multiple cue locations. In order to perform the task, the monkeys now needed to observe and remember the location of the cue, and compare it with the location of the second stimulus in order to determine if that was a match or not and plan the appropriate response. We anticipated that expanding the range of stimulus locations would produce further changes in neural recruitment. Indeed, responses to the cue stimulus, which now became essential for the task, increased greatly (Fig. 3d). However, by virtue of presenting the cue at multiple locations, more neurons had a chance of being activated (see also active units in Fig. 1k) whereas, no such change occurred in the presentation of stimuli in the passive task. Progression of training in this phase was characterized by stability in other aspects of neural activity; no change in baseline firing rate was evident between early and late training phases (Fig. 4f, two-tailed t-test, t (234) = 0.73, p = 0. 468) and these negative findings were also shared in the passive tasks (Fig. 4j, twotailed t-test, t (95) = 0.02, p = 0. 987).

Phase IV amplified the working memory demand of the task, as the duration of each of the two delay periods in the trial progressively increased from 0.25 s to 1.25 s. The most salient change in neural activity was the increase in firing rate during the delay period relative to the baseline (Fig. 3b). As the timing of task events changed for the first time during training, the ramping of activity after the cue presentation also disappeared (Fig. 3d). This change occurred rapidly, as soon as the delay period began increasing in the active task (Fig. S4). The elimination of ramping activity has been previously described in working memory tasks that randomize the delay period compared to versions of the task with a fixed delay period ^35^. As was the case in phase III, some of these changes were transient. The absolute level of activity declined later in the phase (this is evident in Fig. 4a as well). We have recently reported an analogous phenomenon of working memory activity becoming more distributed across a larger population of neurons, while individual activity decreases, in an experiment relying on single-neuron recordings at early and late phases of a working memory task with multiple stimuli ^26^. Increasing the delay period of the active task also induced long-lasting changes in the prefrontal network, which were evident in recordings during the passive task: Increased delay period relative to baseline was now evident in passive recordings, the only phase in which this occurred (Fig. 3e, two-tailed t-test, t (67) = 7.17, p = 7.49 × 10^-10^). A decrease in the baseline firing rate was also observed in the passive task (Fig. 3f).

In addition to analyzing responses to the best location of each neuron in the passive task based on the phases of task learning, it was also important to examine how responses to the same location changed as a function of experiencing these stimuli in the context of the task. The first two phases of the active task involved training with stimuli always presented at the same two locations, in the left and right of the screen followed by choice targets at orthogonal locations, at the top and bottom. Responses to stimuli at other locations in the passive task were altered during this period even though the monkey had not actively been trained with them yet. Such an example change from the passive to the “pre-choice” stage is shown for the lower-right location in Fig. S5. A 1-way ANOVA test indicated a significantly different firing rate at the four training phases (F_3,223_ = 5.89, p = 6.97 × 10^-4^). In the middle of phase III, the lower-right location became the site of one of the two choice targets in the active task, when the cue and match stimuli were first presented in the locations diagonal to it, in the upper-right or lower-right location (see Fig. 1d). This was also associated with a large increase in firing rate for the presentation of the stimulus in the lower-right location in the passive task (“choice stage” in Fig. S5). Finally, when the monkey was exposed to stimuli appearing at the lower right location as cues that needed to be remembered in phases III and IV, responses to stimuli at that location actually declined in the passive task (“cue stage” in Fig. S5). These results suggest that transfer of activity changes in response to a stimulus were not tethered to the specific stimulus being used in the context of the active task but were more general, as the network was altered during training.

Although we emphasized changes in neuronal activity during the execution of the passive task, it is important to point out that several other aspects of neuronal responses remained stable in the course of training. We used the demixed Principal Component Analysis ^36^, to formally identify the types of information represented in the activity of the passive task. We have previously documented that major differences characterize the transition from the execution of the passive task in the pre-trained phase to the execution of the active task after training ^36^. We now found that the representation of stimulus locations, match or nonmatch status of a trial, and invariant components remained fairly unchanged in the activity of neurons during the execution of the passive task across the training phases (Fig. S6). Most importantly, decision components, which represent information about the match and nonmatch status of the second stimulus were virtually absent in the passive task across all training stages. The conclusion was confirmed by a decoding analysis (Fig. S7). Although the decoder readily extracted the location of the first stimulus, the match or nonmatch status of the second stimulus could barely be decoded with above chance accuracy from the passive task at any phase of training, in stark contrast with the same information being decoded from the active task (two-tailed t-test, t (5) = 3.92, p = 0.011). These negative findings provide assurance that the changes we did observe in the passive task represent a true transfer of neural effects across tasks, rather than implicit execution of the active working memory task, even during passive fixation.

## Discussion

It has been recently recognized that working memory ability is malleable and can be increased by using computerized training ^4–6^. After such training, performance improvements generalize between tasks by improving not only for the trained tasks but also for tasks that were not part of the training ^5, 8–10, 37–40^. Our study allowed us a window on the changes of the prefrontal circuitry as the result of such training-induced plasticity. Across four learning phases that required mastery of different conceptual elements and induced qualitatively distinct changes in neural activity, we consistently observed that neural changes in the prefrontal network through training in the active task transferred into the passive task. Changes of neuronal activation in the active task included changes in LFP power, MUA responsiveness, and single neuron firing rate, in agreement with changes previously documented in single-electrode studies comparing different populations of neurons, recorded at different training stages ^26, 41–44^, or during the course of a daily training session, when a specific stimulus is associated with reward ^45, 46^. Both increases and decreases in activity observed in the active task transferred to the passive task, as did null results (e.g. no baseline activity change during the course of Phase III). Artificial neural networks have provided a framework for understanding transfer learning: a network trained on one task produces changes in connection weights in the hidden layers of the network, which when probed with a different task generate training-dependent output ^47^. We now document the neural equivalent of this process, as learning takes place.

An important consideration for the interpretation of the findings is whether the effects observed in the passive task were the consequence of monkeys mentally “performing” the active task even when presented with stimuli, passively. This possibility is unlikely for multiple reasons: Blocks of trials of the passive task were presented in exactly the same routine fashion, at the beginning of the session every day. The passive task did not involve target stimuli at the end of the trial, allowing the monkeys to realize that no choice was required, from the first trial of the block. The timing of stimulus presentation differed between passive and active tasks, at least through the first three stages of training (until the duration of the delay period was increased), again making the two tasks appear very different. The first two phases of the active task involved training with stimuli always presented at the same two locations, in the left and right of the screen. Yet, responses in the passive task were altered during this period even for stimuli that the monkey had not actively been trained with yet. Nor was the monkey able to easily generalize performance of the active task with stimuli appearing at other locations; the entire duration of phase III training was devoted precisely to this purpose. Information about the match or nonmatch status of stimuli, on which decisions are based, and which differs in correct and error trials of active working memory tasks ^48, 49^, was also minimal in the passive task.

Working memory is thought to be mediated by persistent activity generated during the delay interval of working memory tasks, though this has been a matter of debate ^27, 28, 50–55^. Our results suggest enduring changes in the prefrontal circuitry after training, increasing its excitability of prefrontal neurons and the ability to generate persistent activity. Alternative models emphasize instead increase of gamma power at times of active memory maintenance ^29–31^, consistent with evidence from EEG studies in humans, which most often associate working memory maintenance with increased gamma power ^56^. However, an increase power in high beta and low gamma frequency, e.g. in the 24-60 Hz range has also been reported in working memory tasks ^57–59^. Guided by these models, we tested for systematic changes in LFP power, and we indeed found consistent decreases in high beta – low gamma power at successive stages of training. Regardless of the underlying mechanisms that brought about these changes at the level of beta-frequency LFP power, these also transferred to the passive task. A salient effect of training was that when probed with passively presented stimuli, larger populations of prefrontal neurons were shown to respond, and to be capable of generating persistent activity in the “delay” period of the task, even though it was not necessary to maintain these stimuli in memory for the requirements of the passive task. Such changes would also be expected to strengthen neuronal responses to other tasks that rely on maintenance of information in mind. Our results provide a framework for probing such changes in future studies.

## Methods

### Subjects

Two male, rhesus monkeys (*Macaca mulatta*) weighing 8-9 kg were used in this study. All experimental procedures followed guidelines by the U.S. *Public Health Service Policy on Humane Care and Use of Laboratory Animals* and the National Research Council’s *Guide for the Care and Use of Laboratory Animals* and were reviewed and approved by the Wake Forest University Institutional Animal Care and Use Committee.

### Surgery and neurophysiology

The monkeys were initially acclimated with the laboratory and trained to maintain fixation on a white dot while visual stimuli appeared on the screen. After this initial stage of training was complete, the monkeys were implanted with a chronic array of electrodes in their lateral prefrontal cortex. The implant comprised an 8 × 8 grid of electrodes, with adjacent electrodes spaced 0.75 mm apart from each other, thus covering an area of 5.25 mm × 5.25 mm. The electrode array targeted the dlPFC, with electrode tracks descending in both banks of the principal sulcus (Fig. 2a). The position of the array was determined based on magnetic resonance imaging (MRI) and verified during the implantation surgery. Electrode depths were adjustable and were repeatedly adjusted to optimize placements, over a period of several weeks. Once electrode positioning was finalized, task training and neurophysiological recordings from the array commenced. Neuronal data from each electrode were recorded throughout the training. Multi-unit data were collected from each electrode from areas 8a and 46 of the dlPFC, using an unbiased spike selection procedure. The threshold for spike acquisition was set at 3.5 × RMS of the baseline signal, for each electrode, each day. The electrical signal from each electrode was amplified, band-pass filtered between 500 Hz and 8 kHz, and recorded and sampled at 30 kHz using a Cerberus system (Blackrock Microsystems, Salt Lake City, UT).

### Behavioral tasks

The monkeys faced a computer monitor 60 cm away in a dark room with their head fixed, as described in detail previously ^43^. Visual stimuli display, monitoring of eye position, and the synchronization of stimuli with neurophysiological data were performed with in-house software ^60^ implemented in the MATLAB environment (Mathworks, Natick, MA), and utilizing the psychophysics toolbox ^61^.

The monkeys were trained in a Match/Nonmatch task involving four phases. The monkeys were then trained to perform a spatial working memory task, requiring them to maintain fixation, observe two stimuli appearing in sequence separated by delay periods, and to indicate if the two stimuli appeared at the same location or not by making an eye movement to one of two choice targets (Fig. 1). The training could be broken down into four phases. The first phase of training involved training the monkeys to make an eye movement to one of two choice targets and determining that only one of them is rewarded (Fig. 1b). The phase began with the monkeys being exposed to match trials, requiring an eye movement to the “Diamond” choice target. The first stimulus appeared always at the same location (to the right of fixation), followed by a very brief delay period activity (0.25s) and a second presentation of the stimulus at the same location. After the second delay period, the two choice targets appeared with the fixation point turning off, either above or below the fixation point, but randomly switching between trials. In the absence of the fixation target, the monkeys quickly foveated one of the choice targets, and they learned through trial and error that the “Diamond” choice target was rewarded. In a subsequent training day, nonmatch trials were introduced. Now the first stimulus appeared again at the right location, but it was followed by a nonmatch stimulus. When the choice targets appeared at the end of the trial, it was the “H” shape that was rewarded. The monkeys quickly reversed and saccaded to the “H” choice target. Phase I of training involved delivering match and nonmatch trials in blocks with decreasing numbers of trials before alternating.

Phase II involved randomly interleaving match and nonmatch trials (Fig. 1c). Through this process, the monkeys eventually associated the concept of “match” with the “Diamond” and “nonmatch” with the “H” shape. Phase II concluded when the monkeys were able to perform the task at 75% correct. This was the most challenging phase of training.

So far in training, the cue stimulus always appeared at the same location. Phase III involved the generalization of stimulus location (Fig. 1d). The first stimulus appeared at a different, followed by a second stimulus at the same location, or its diametric. Choice targets appeared orthogonal to the axis defined by these possible stimulus locations. To facilitate learning, whenever a new location was introduced, we relied again on blocks of match and nonmatch trials. To ensure that the monkeys did not “forget” the previous location, they continued to practice these, and every time a new location was added, randomized trials involving all trained locations were interleaved together. The monkeys were able to progress much faster through this stage, though they did not automatically generalize when a new location was introduced. Some practice was necessary to determine what the appropriate choice was for match and nonmatch stimuli appearing at these novel locations.

The final phase of training, Phase IV, involved progressively increasing the delay period duration. Both delays period between the first and second stimulus, and between the second stimulus and choice targets increased in tandem. Durations varied from 0.25 s to 1.5 s.

At the onset of the working memory task training, the monkeys were already able to maintain fixation, and had already been exposed to the visual stimuli that would eventually be incorporated in the task (white squares, appearing at one of nine locations). The timing of the stimulus presentation mirrored the final phase of the task (Fig. 1a). The only difference was that the choice stimuli were presented at the end of the trial, and the monkeys were rewarded for maintaining fixation after the second delay period. An initial set of recordings was obtained from the chronic array at this phase, providing a baseline of neuronal activity prior to the task training. Additionally, the passive presentation of stimuli continued throughout training; the first block of trials presented every day involved the exact same passive stimulus presentation. Thus, monkeys were aware that they did not need to perform a working memory task.

### LFP Analysis

We used the FieldTrip toolbox ^62^ for preprocessing analysis and the Chronux package ^63^ for time-frequency analysis. A bandpass filter (0.5-200 Hz) was first used. We removed line power (60 Hz) from each electrode and trial, if present. We used a generalized linear model to identify electrodes with variance outliers, and we omitted from the analysis. Therefore the number of electrodes that were averaged varied from 45~60 in each trial. We then used a multi-taper method to perform a power spectrum analysis of LFP. Power spectra were constructed from all trials and electrodes in each session and then averaged across sessions after subtracting the mean power of the baseline fixation period at each frequency. We then compared the LFP power at each frequency between the control and simulation conditions. We also analyzed the LFP power at different frequency bands defined as alpha (8-14 Hz), beta (20-45 Hz) and gamma (46-70 Hz). Line-plots were constructed based on average and standard deviation across sessions (treating one session as one observation). One-way ANOVA was used to compare LFP power between phases, at each frequency band.

### Spiking Data Analysis

All data analysis was implemented with the MATLAB computational environment (Mathworks 2019, Natick, MA). We identified MUAs that were responsive to the task and informative about the stimuli as those whose mean firing rate to the different stimulus conditions were significantly different from each other, determined by 1-way ANOVA (P < 0.05). The ANOVA was performed for the firing rate averaged across the entire cue period, and the delay period and compared across available cue locations (typically 9). For task conditions that involved only one cue location (active task, Phase I and II), responsive neurons were identified as those with firing significantly exceeding the fixation period firing rate (paired t-test, P < 0.05) between either the first stimulus presentation or the delay period. We additionally required a minimum 10% firing rate increase during the stimulus presentation over the fixation interval, to avoid false positives. Responsive single neurons were determined in the same way as MUA, except without requiring the 10% proportional increase of activity. Firing rate analyses presented here relied on data from correct trials. For each neuron, we identified the cue location that elicited the best response during the cue presentation period, and during the delay period, determined independently. Activity of the best location in each day, which was defined by the maximum activity in cue or delay periods, was shown in heat maps. Daily responses were evaluated by calculating the average firing rate among all selective sites recorded. To compare active and passive conditions, data from responsive neurons recorded in the active and passive conditions were plotted.

Decoding analysis was carried on the stimulus direction and decision type (i.e., match or nonmatch) factors. Therefore, the chance performance for stimulus direction decoding was 12.5 %, for decision type decoding was 50%. The analysis was carried out using leave one trial out cross-validation. The model was fit with the remaining trials and tested on the trial that was held out of the analysis. The decoding accuracy of each neuron population was computed in 200 ms bins, advanced in 20 ms increments. In order to facilitate comparison, the same number of neurons (50 neurons for the passive task, and 100 neurons for the active task) were used across different training phases with 50 times of repetition. The posterior probability of stimulus or choice, which is the probability of selection for the stimulus location or selecting the decision type over trials was calculated by:

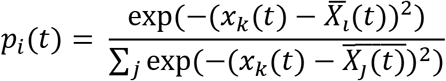

Here *p_i_*(*t*) represents the probability for option *i* at time t, *x_k_*(*t*) represents the neural population activity in a single trial *k*, at time t. The variable 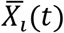 represent the mean neural population activity across trials with specific stimulus location or decision type, which were indicated by *i* or *j*.

Demixed Principal Component Analysis (dPCA) was performed as we have described elsewhere ^36^. This method decomposes population activity into the stimulus components (8 peripheral stimulus locations, excluding the foveal location) and the decision components (match or nonmatch). The method treats the responses of each neuron to one type of stimulus condition as one dimension and then performs dimensionality reduction to determine components that correspond to stimulus and task variables.

## Data availability

All data will be posted to Github or made available upon reasonable request https://github.com/ChristosLab

## Code availability

All code will be posted to Github or made available upon reasonable request https://github.com/ChristosLab

## Acknowledgments

Research reported in this paper was supported by the National Eye Institute of the National Institutes of Health under award number R01 EY017077 to C.C.; by NINDS training grant T32 NS073553; NIMH grant F31 MH104012 to M. R. R.; and by the Tab Williams Family Endowment. We wish to thank Kathini Palaninathan, Aquil Jones, Austin Lodish, Leonardo Silenzi, Rafael Mendoza, Macrae Robertson, and Mia Allen for technical help.

## Contributions

C.C., M.R.R. and D.T.B. conceived and designed the experiments. M.R.R., H.T. performed the experiments. H.T., M.R.R., X.L.Q., B.S. and C.C. performed data analysis. C.C. wrote the manuscript with input from all authors.

## Corresponding Author

Correspondence to

## Ethics declarations

The authors declare no competing interest

## Supplementary Material

### Supplementary Results

#### LFP Analysis

We focused on differences in LFP power differences in alpha and beta frequency bands during the cue presentation period, because they were most characteristic of changes that occurred with training, but these were not the only changes evident. Here we present results from additional task periods. A significant difference in beta-frequency power between phases (1-way ANOVA, F_3,889_ = 96.3, p = 5.54 × 10^-54^) and alpha-frequency power (F_3,889_ = 197.7, p = 3.22 × 10^-98^) was observed over the fixation period. Similar changes occurred during the passive task, with a decrease in beta-frequency power (F_4,446_ = 34.8, p = 2.49 × 10^-25^) as training progressed and a significant increase in alpha-frequency power (F_4,446_ = 23.1, p = 2.31 × 10^-17^). During the cue presentation period, we observed significant difference beta-frequency power (F_3,889_ = 106.4, p = 7.05 × 10^-59^) and alpha-frequency power (F_3,889_ = 99.5, p = 1.55 × 10^-55^) in the active task; beta-frequency power (F_4,446_ = 19.2, p = 1.36 × 10^-14^) and alpha-frequency power (F_4,446_ = 16.4, p = 1.37 × 10^-12^) in passive task respectively. We also observed a significant difference in beta frequency power (F_4,446_ = 28.7, p = 2.89 × 10^-21^) and (F_4,446_ = 22, p = 1.31 × 10^-16^) during the first and second delay period in the passive task. However, there was no significant difference in alpha-frequency power in delay period 1. As training progress there was a slight increase in alpha-frequency power (F_4,446_ = 3.9, p = 0.004) at delay period 2.

### Supplementary figures and tables

**Fig. S1.**
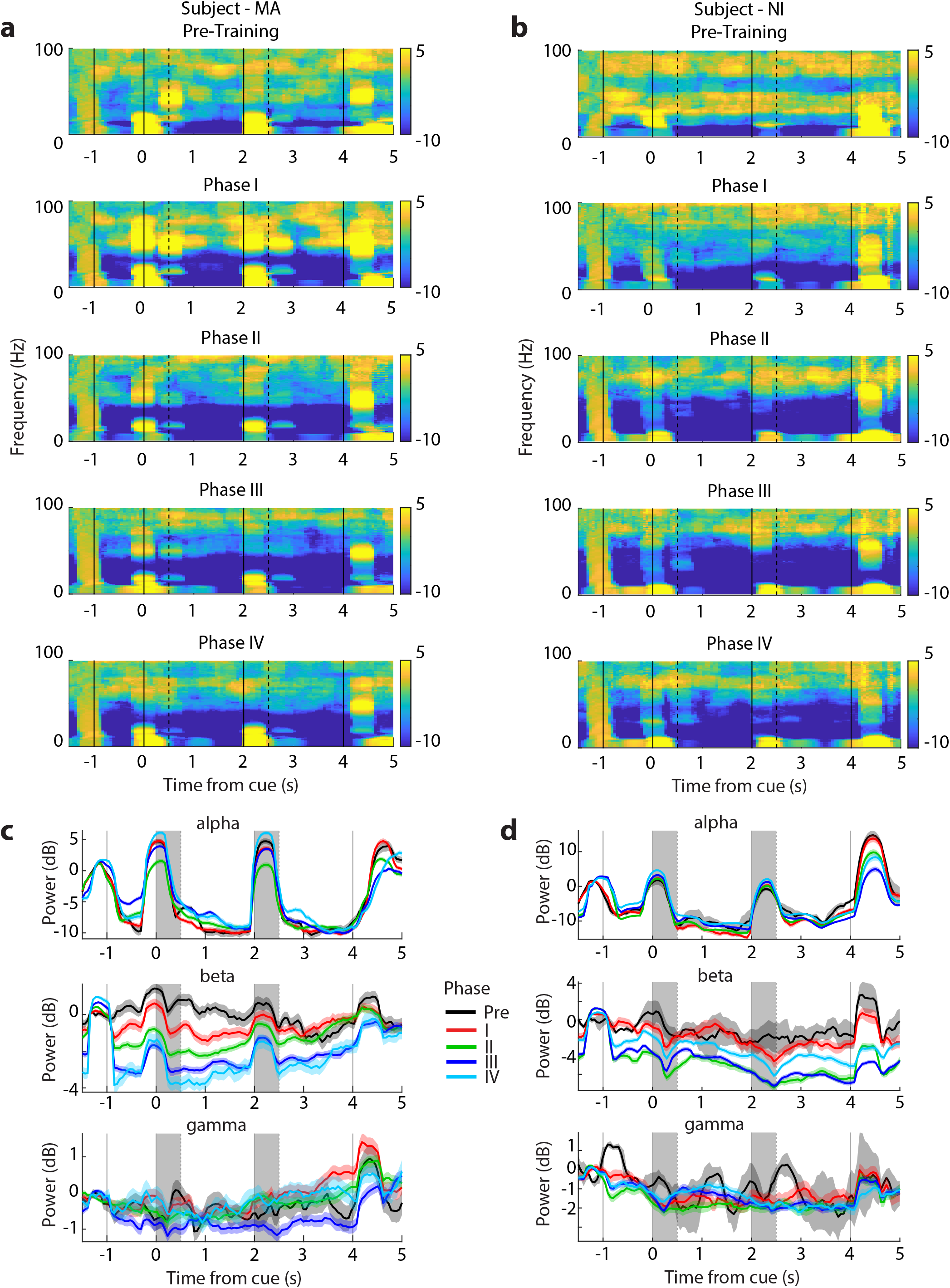
LFP analysis in two monkeys. **(a)** LFP power spectrum as a function of time for the training phases in the passive task from monkey MA. **(b)** LFP power spectrum as a function of time for the passive task, as training progressed in the active task. Time course of power at discrete frequency bands and different training phases of the active task: alpha (8-14 Hz), beta (20-45 Hz), gamma (46-70 Hz). **(c-d)** As in a-b for the second monkey subject NI.

**Fig. S2.**
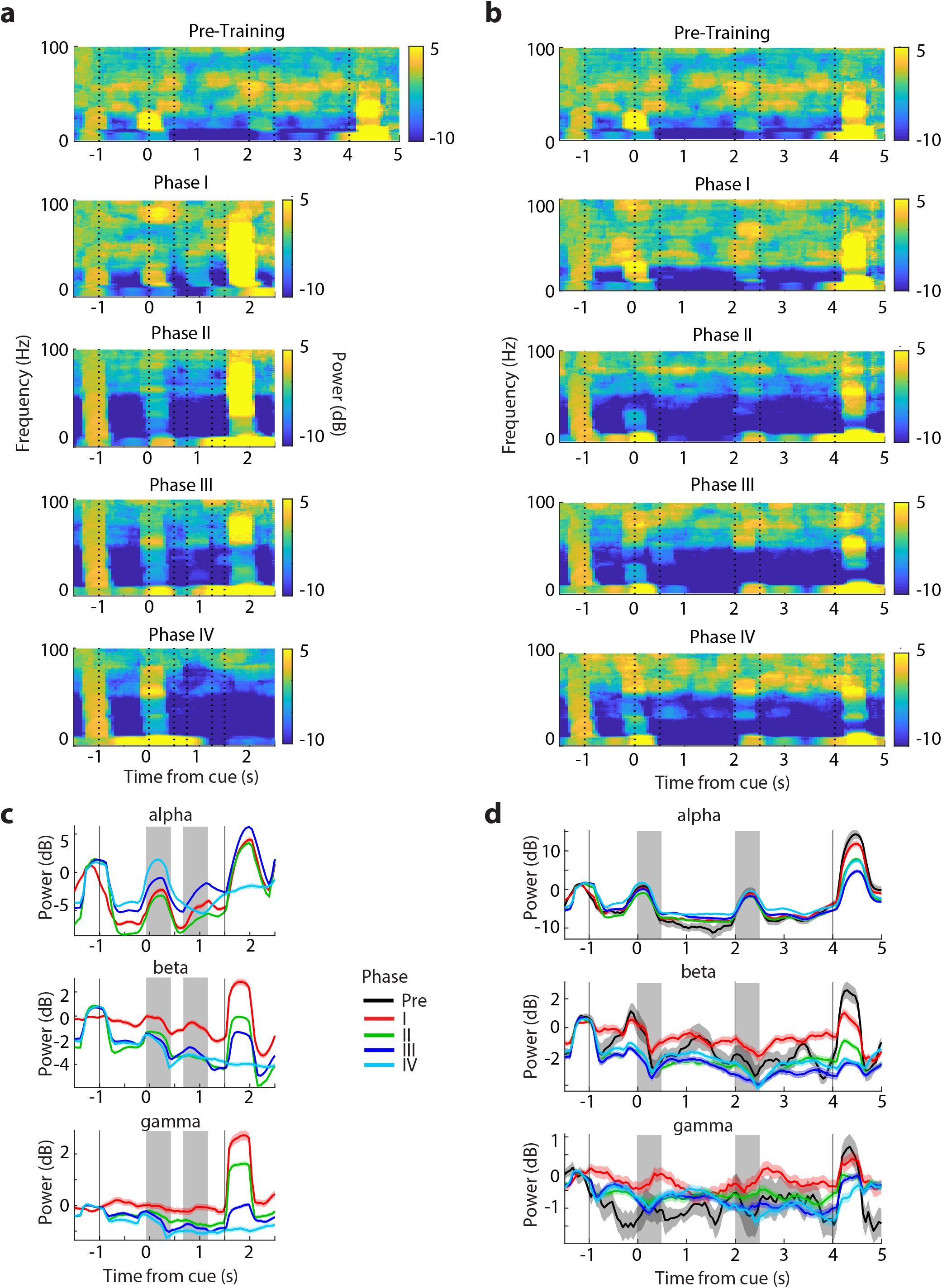
LFP analysis in electrodes that yielded single units. LFP power spectrum as a function of time for the training phases in the active **(a)** and passive **(b)** tasks constructed only from electrodes that yielded single neurons (ensuring therefore that recordings were still active). Conventions are the same as Figure 2.

**Fig. S3.**
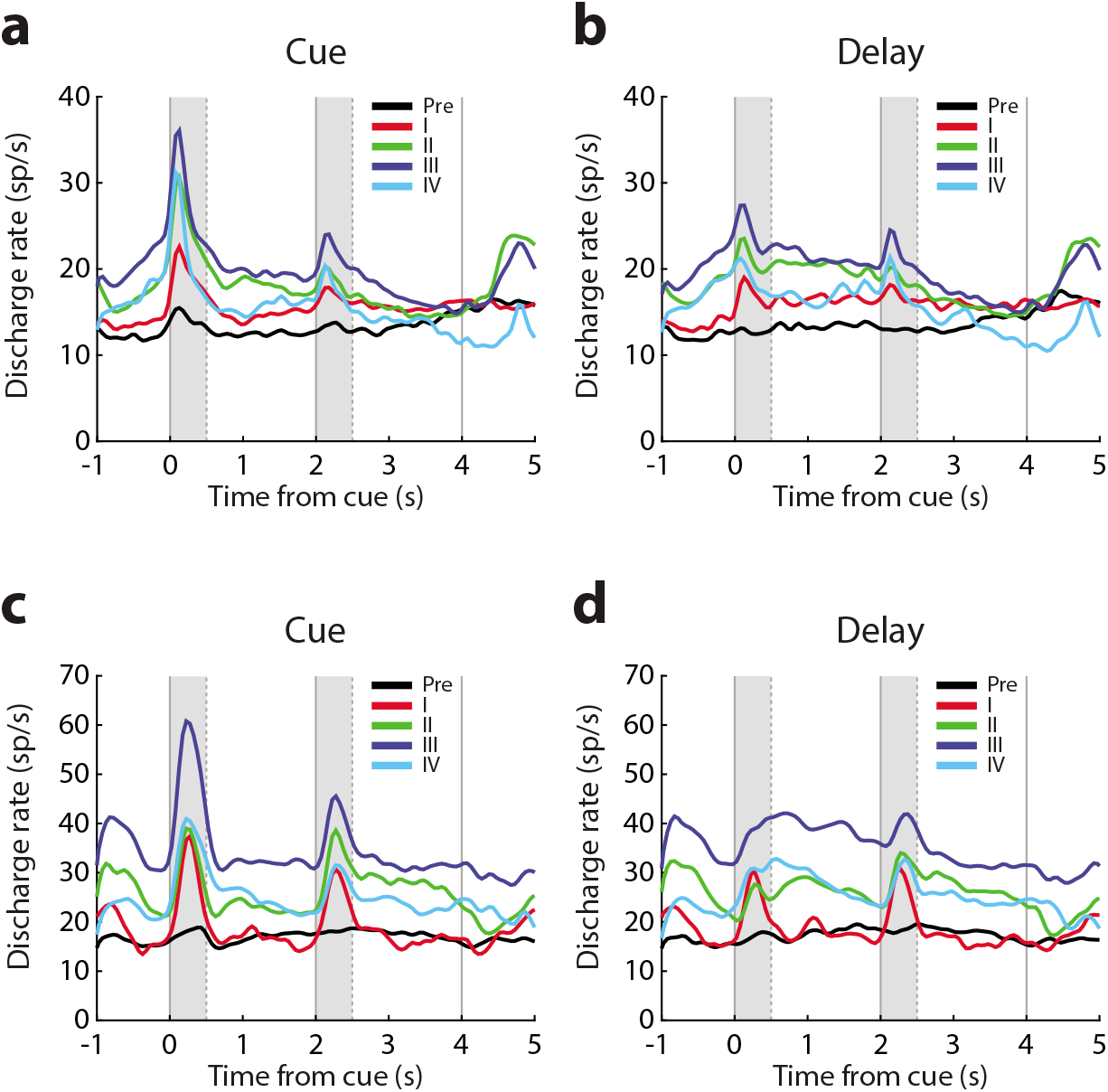
MUA changes in activity across single channels. **(a-b)** Population PSTH constructed based on responsive MUA units of the passive task, identified in each training phase. Data have been selected based on the presentation of the best cue (a) and delay period (b) activity, from responsive MUA units always isolated from the same electrode of subject MA (n = 324). **(c-d)** As in a-b for the second monkey subject NI (n = 213).

**Fig. S4.**
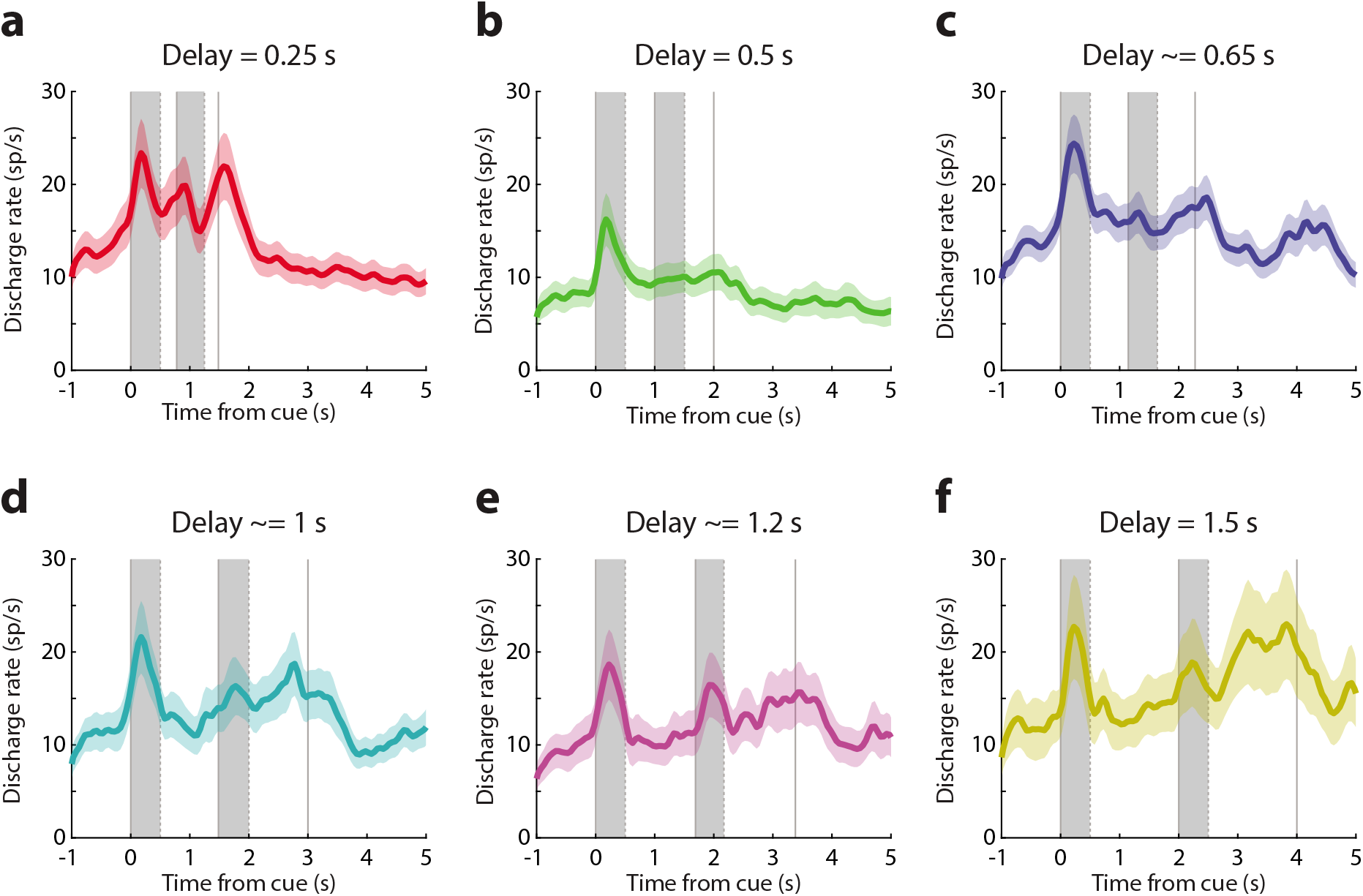
Single neuron responses during the increase of the delay period. Population PSTH is drawn for the activity of responsive single neurons at different subphases of Phase IV of training, when the delay period of the task was progressively elongated. Details of the changing of delay length can be found in Figure 4a. Shaded zones represent mean ± SEM. Cue and match presentations are indicated with gray bars.

**Fig. S5.**
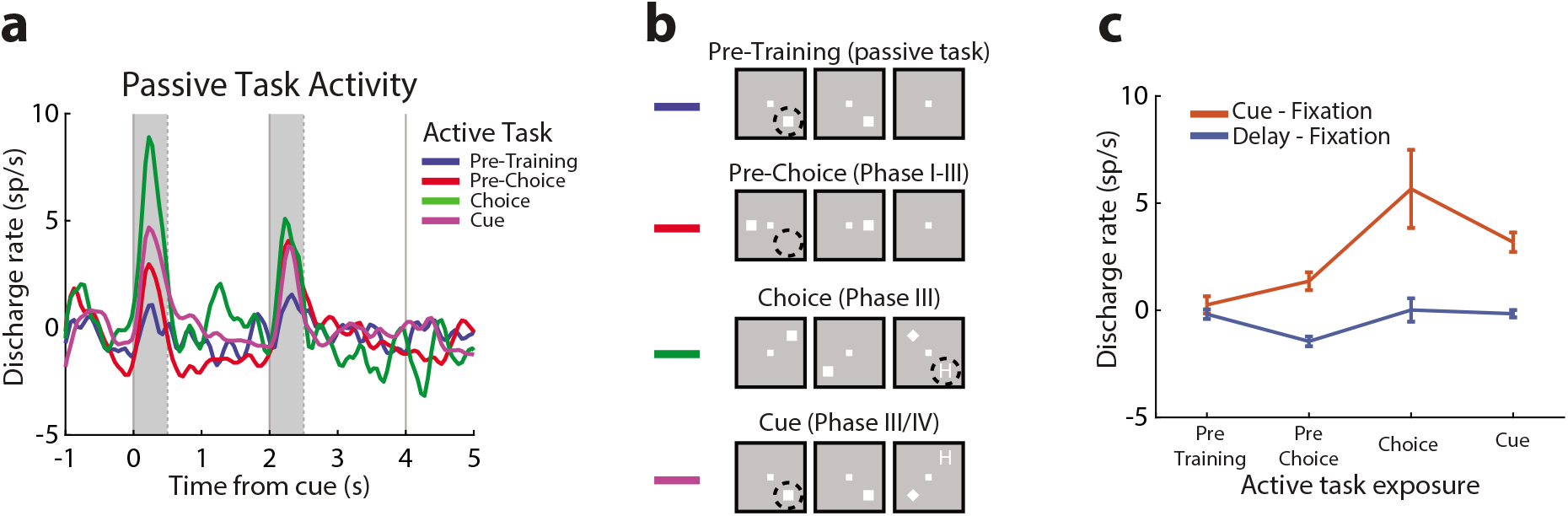
Response to the same location across training. **(a)** Mean activity of single neurons in the passive task, responding to the lower-right location, as a function of time, now grouped based on exposure to stimuli appearing in this location in the context of the active task. **(b)** Pre-training phase refers to the period before active training began. “Pre-choice” refers to the period between the beginning of active training and the first time that any stimulus appeared in this location in the active task. The “Choice” period begins the first time that a choice target stimulus appeared in this location, during phase III, when the cue and nonmatch stimuli appeared at diagonal locations. The “Cue” period begins the first time that a cue stimulus appeared at that location. **(c)** Mean evoked firing rate of units responded to the lower-right during the cue and delay period (n = 903).

**Fig. S6.**
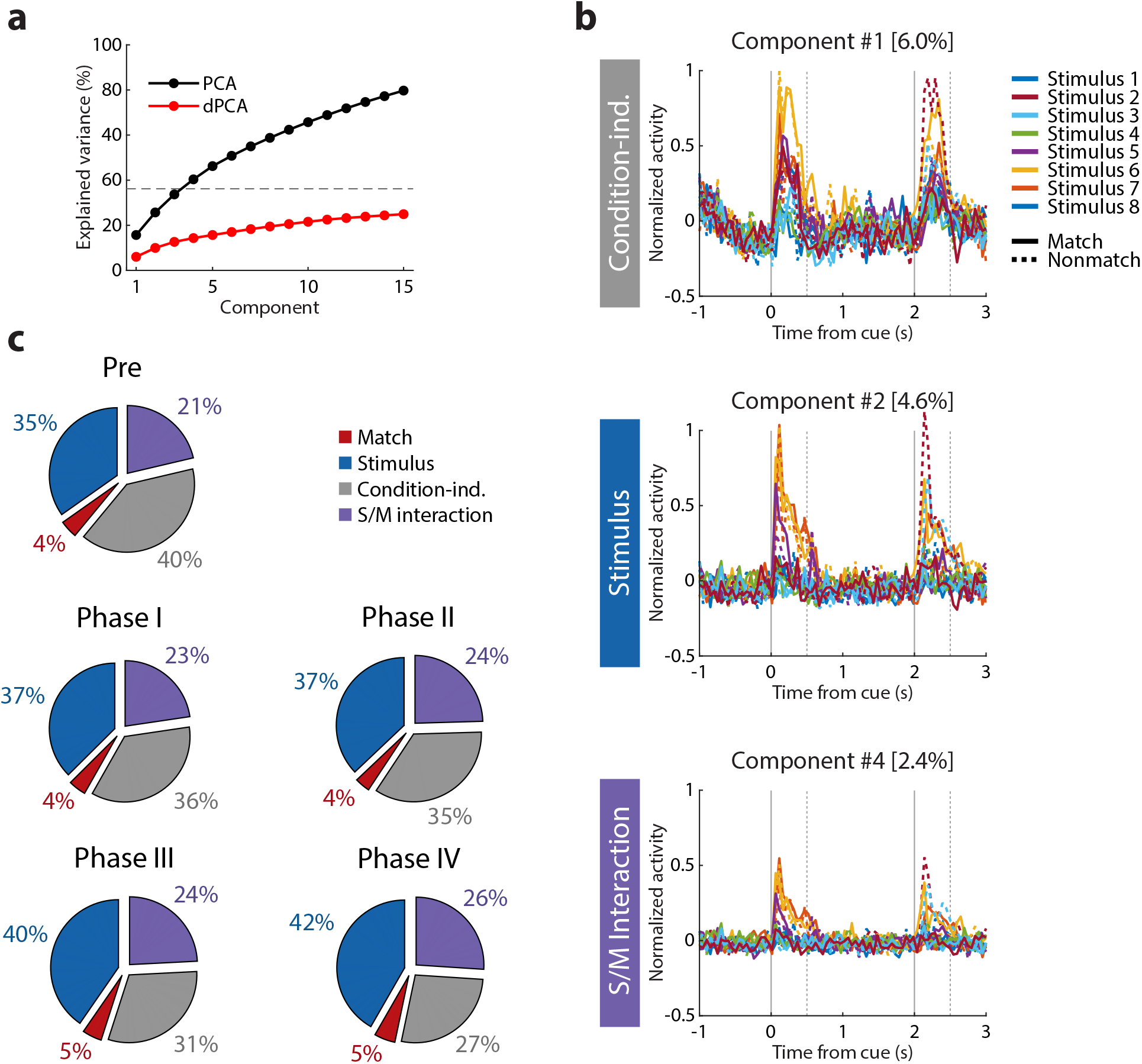
Demixed Principle Component Analysis. **(a)** Cumulative variance explained by PCA (black) and dPCA (red) for the passive task in Phase II. Dashed line shows an estimate of the fraction of “signal variance” in the data. **(b)** Three components of dPCA analysis based on results of Phase II: a stimulus-related component, a stimulus/decision mixture, and a condition independent component. **(c)** Pie charts represent the percentage of variance explained by each type of component (stimulus location, the match or nonmatch status of the trial, condition-independent components, and mixtures thereof) in the responses of single neurons during the passive task, across training stages.

**Fig. S7.**
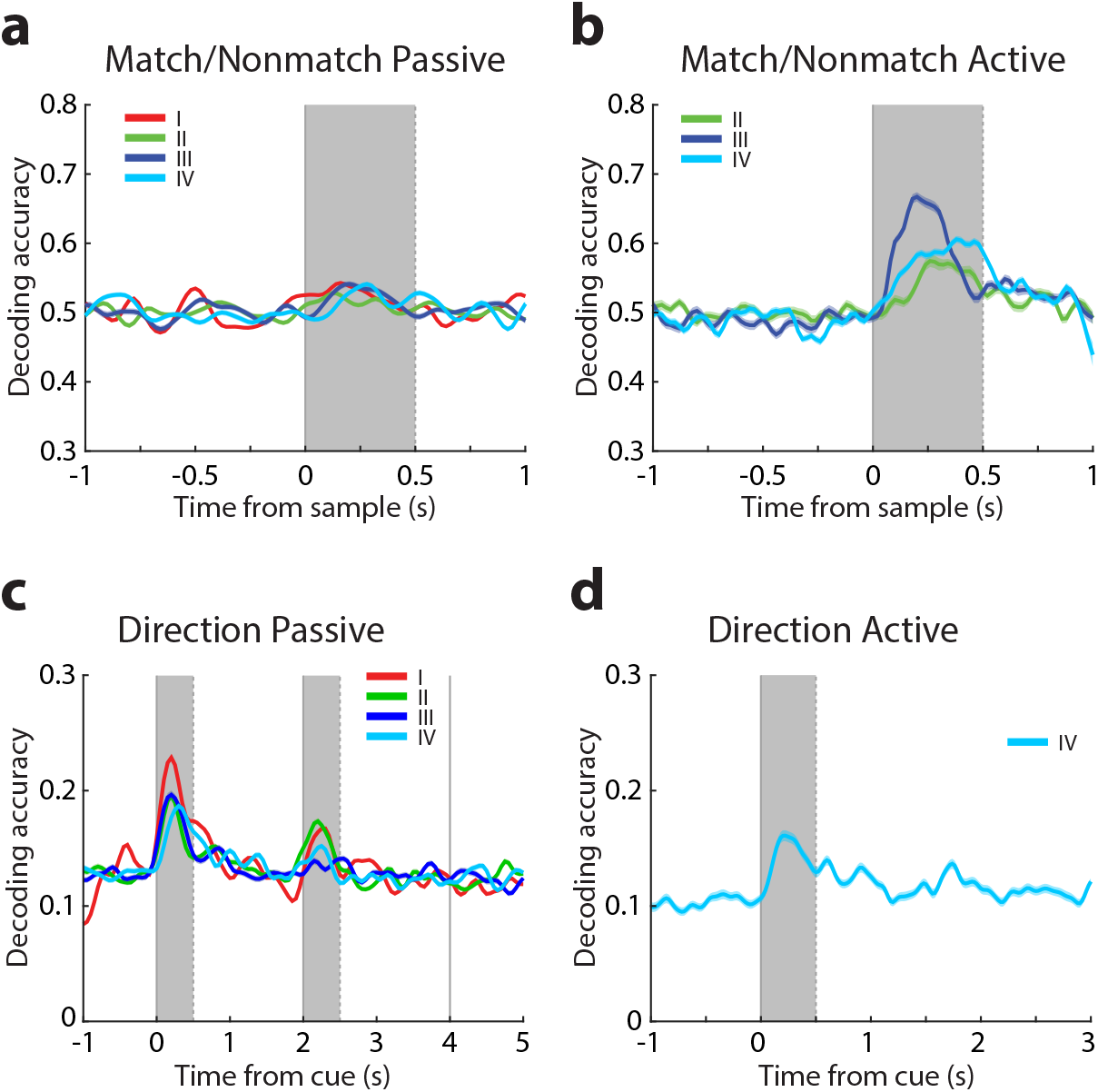
Decoder Analysis. **(a-b)** Accuracy of decoding the match or nonmatch status of the second stimulus based on single neuron responses pooled from the passive task (a) the active task (b), plotted separately for each training phase. Only phases II-IV are included in the active task, as only in these the monkey has been trained to distinguish between match and nonmatch choices. **(c-d)** Accuracy of decoding stimulus locations based on single neuron responses pooled from the passive task (c), and the active task (d), separately for each training phase. Only phase IV is included in the active task, as only in this the monkey has been trained to distinguish all locations.

